# A Low-Cost Stage-Top Incubation Device For Human Cell Imaging Using Rapid Prototyping Methods

**DOI:** 10.1101/2023.09.29.560179

**Authors:** Michael Worcester, Melissa Gomez, Pratyasha Mishra, Quintin Meyers, Thomas E. Kuhlman

## Abstract

Live imaging of human or other mammalian cells at multi-hour time scales with minimal perturbation to their growth state requires that the specimen’s optimal growth conditions are met while fixed to a microscope stage. In general, ideal conditions include culturing in complete growth media, an ambient temperature of 36-37 C, and a humidity-controlled atmosphere comprising typically 5-7% CO_2_. Commercially available devices that achieve these conditions are not a financially viable option for many labs, with the price ranging anywhere from $12000 to $40000. The advent of 3D printing technology has allowed for low-cost rapid prototyping with precision comparable to traditional fabrication methods, opening the possibility for in-lab design and production of otherwise prohibitively expensive equipment such as stage-top incubation devices. The continued usefulness and widespread availability of single-board computers (SBC) such as Arduino and Raspberry Pi also simplify the process by which these devices can be controlled. Here we report the production of a do-it-yourself (DIY) device for stage-top incubation with temperature and atmospheric control with a cost reduction of approximately 100x.

## I. INTRODUCTION

Human and mammalian cell health *in vitro* is maintained partly through use of media supplemented with a bicarbonate buffering system. Environmental CO_2_ at concentrations typically ranging from 5-7% dissolves into the medium and allows for regulation of internal pH usually between 7.2 and 7.4 [1]. Maintenance of these optimal growth conditions in the lab is typically accomplished via incubators that actively pump CO_2_ into the interior. Generally, laboratory incubators are not conducive to facilitating the use of a microscope, making it infeasible to perform frequent microscopy of samples without perturbing the cells unless an environmental chamber for the microscope is purchased or assembled, typically at great financial or labor cost.

The geometry of an inverted microscope, where the objective lens sits underneath the sample, is advantageous for adherent cell cultures, where individual cells settle to the bottom of wells with the assistance of gravity [2]. This arrangement also makes it feasible to use an incubation device sitting on the top surface of the stage to maintain ideal environmental conditions for imaging live human cell cultures over long periods with minimal perturbations. Imaging of the sample with an inverted microscope means that any culturing device must have an open window on the bottom face.

Commercially available stage-top incubation devices create a sealed atmospheric environment for the sample that can be quickly accessed and regulated with a series of sensors and controllers [3]. Typically, caution is not taken to make the device airtight, and instead the system opts for a constant flow of mixed air-CO_2_ atmosphere in and out of the device, with humidity control to reduce evaporation from the sample. Purchasing of commercially available devices can cost between approximately $12,000 to $40,000 in 2023, depending upon the features and manufacturer of the device. Such high cost is a significant barrier of entry for many laboratories. Lower cost custom solutions for mammalian cell incubation reported in the literature are generally suitable only for custom built microscopes and stage arrangements and are typically not “plug and play” stage top devices [4, 5]. Here, we describe a DIY stage-top incubation device, we call the “DIYncubator”, that can be constructed simply for approximately two orders of magnitude lower cost than commercial systems (*∼* $250 USD) and is compatible with existing commercial or custom microscopes and stages.

## II. METHODS

### A. DIYNcubator Design and Assembly

The chassis of the DIYncubator (Fig. 1) was designed in Autodesk Inventor 2022, formatted for 3D printing with Ultimaker Cura software, and printed on an Ultimaker S7 using black acrylonitrile butadiene styrene (ABS) plastic. Machine screws and threaded screw inserts were used to attach the lid to the device. The contact region between the two major halves of the chassis was lined with neoprene to make the device approximately airtight, limiting the amount of gas required to maintain a constant atmosphere. It is not necessary to make the device completely airtight, but reduction of leakage helps reduce the amount of CO_2_ necessary and therefore the cost of operation. Attachment and sealing of other permanent components of the device was achieved with polyurethane or cyanoacrylate adhesive and sealed with silicone caulk typically used for kitchen and bath projects. The shape of the DIYncubator was designed to attach to a Prior ProScan III stage on a Nikon Ti-E inverted microscope. Minor design alterations may be necessary when using a different stage design, but are straightforward to accomplish using 3D modeling software.

**FIG. 1.**
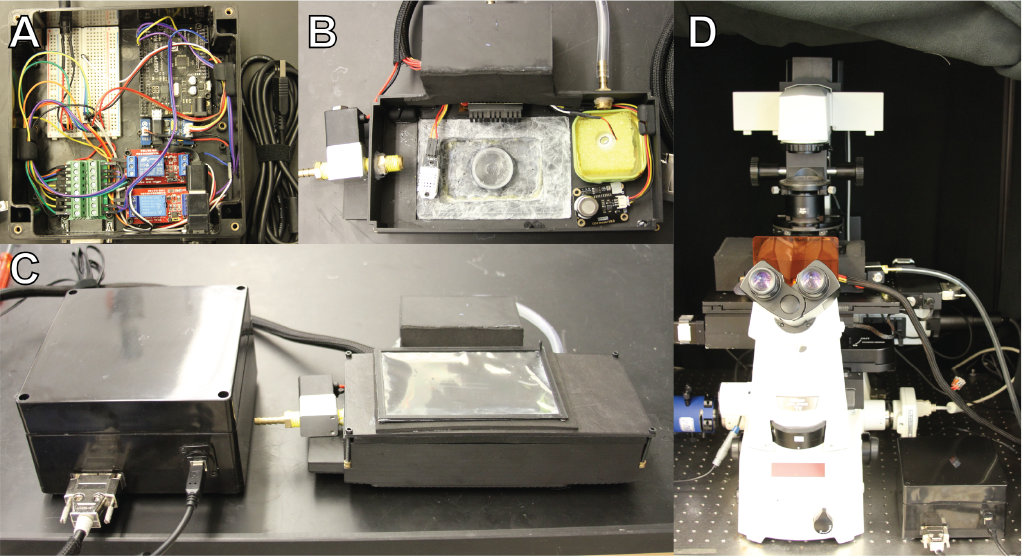
(A) The electronics box, containing the Arduino, relays driving the CO_2_ solenoid and heating unit, atomizer control board, a small breadboard for connections, and a female DB15 port. (B) Photograph of the bottom half of the DIYncubator chassis with electric components including heating unit and fan, CO_2_ solenoid, AM2302 humidity and temperature sensor, MG-811 CO_2_ sensor, and ultrasonic atomizer with water reservoir. Note that the bottom of the device and the adapter for holding the sample (here a 29 mm circular dish) is coated with Parafilm to seal the adapter into the chassis and limit movement of the sample. (C) Assembled electronics box and DIYncubator. (D) The DIYncubator deployed, with insulating neoprene cover, on our Nikon TiE inverted microscope.

The top half of the chassis was designed with a large window to provide quick access to the sample. A covering for the window was constructed from a 3D printed ABS black plastic frame, thin transparent plastic recovered from recycled packaging, and refrigerator magnets for easy detachment from the chassis. The borders of this window were also lined with neoprene to aid in maintaining an airtight environment. The bottom half of the chassis contains a large opening into which a variety of adapters for holding differently sized and shaped samples can be placed. We then coat both the bottom of the chassis and the adapter in Parafilm, as seen in Fig. 1B, which serves both to seal the adapter while in place and also limit movement of the sample when placed on the adapter. Finally, the heating system is placed within a housing at the back of the chassis, and is then covered with a layer of neoprene. To aid in insulation and temperature control of the system, we also fashioned a rough neoprene cover for the entire unit out of scraps remaining from sealing the other components, seen in Fig. 1D.

### B. Electronics, Environmental Measurement, and Control

The control system of the DIYncubator was achieved with an Arduino Uno SBC. CO_2_ and temperature/humidity readings were taken with a MG-811 CO_2_ sensor and an AM2302 temperature and humidity sensor respectively. Alternatively, a K30 CO_2_ sensor can be used, which will be more robust in the long term and provide protection against high humidity. Heat for the system was generated with two flexible polyimide heating film units covered with heatsinks and distributed with a small fan. During operation, the CO_2_ and temperature/humidity sensors were placed as closely as possible to the sample. Gas intake was regulated by a solenoid valve connected to a pure CO_2_ gas tank. Humidity control is provided by an NGW-1pc ultrasonic water atomization unit. As a water reservoir for the atomizer, we used an empty mint tin from Trader Joe’s that holds 45 - 50 mL of water.

The Arduino was programmed to implement a very simple “bang-bang” control scheme for all parameters, where when the measured value of the parameter fell below/above a set point, the corresponding control device was turned on/off with 100% power. If more precision is required, more complex (and expensive) PID control schemes employing MOSFETs could be implemented. The device was interfaced with a desktop laboratory computer via USB connection, and an executable program for user control of the system was developed in Microsoft Visual Studios 2019 for Windows operating systems. The executable program is not absolutely necessary for the device and could be circumvented using the Arduino with an integrated LED display and buttons for operation.

A parts list, design files, and software are available at https://kuhlmanlab.ucr.edu/downloads/.

### C. Human Cell Culture

HEK-293T cells were cultured in complete media containing 90% DMEM with high glucose (Gibco), 10% heat-inactivated FBS (Gibco), 200 mM L-Glutamine (100X), 10 mM MEM Non-Essential Amino Acids (100X), 100 mM MEM Sodium-Pyruvate (100X), and conc. Puromycin (Gibco). Seeding was done at 1×10^4^ cells/cm^2^ in 29 mm glass cover slip dishes (Cellvis) and cells were incubated at 37°C and 5% CO2 for 24 hours prior to imaging.

### D. Sample Microscopy and Analysis

Cells were imaged using a Nikon Ti-E Inverted Microscope and with stage-top incubation either with the DIYncubator or an Ibidi Silver Line Stage Top Incubator. Fluorescent proteins were excited with a Nikon Intensilight Epi-fluorescent Illuminator and imaged every 10 minutes. Images were analyzed using ImageJ [6], CellProfiler [7], and using a custom Python script.

## III. RESULTS

### A. The DIYncubator Provides Stable Environmental Control Over Long Periods

The DIYncubator was constructed for a total cost of*∼* $250. Some additional but not required components, such as a plastic project box to house the electronics, DB15 connectors to simplify connections and deployment, and cable cover and management sleeve brought the cost to *∼* $300. It should be noted that purchasing of materials is up to the user’s discretion, and more careful sourcing or building custom circuitry may lead to further reduction in cost. For example, some of the materials used were recycled from previous experiments and shipping packaging. Creativity is encouraged in building such a device, provided that the solution is still functional.

Using this feedback and control system (Fig. 2), internal CO_2_ levels, temperature, and humidity are stably maintained over long periods (Fig. 3), limited primarily by the volume of the water container used by the atomizer to supply humidity, which we have found to maintain humidity for up to *∼* 20 hours. However, the removable lid of the box provides quick access to easily refill the container with minimal perturbation to the sample, allowing for maintenance of conditions essentially indefinitely.

**FIG. 2.**
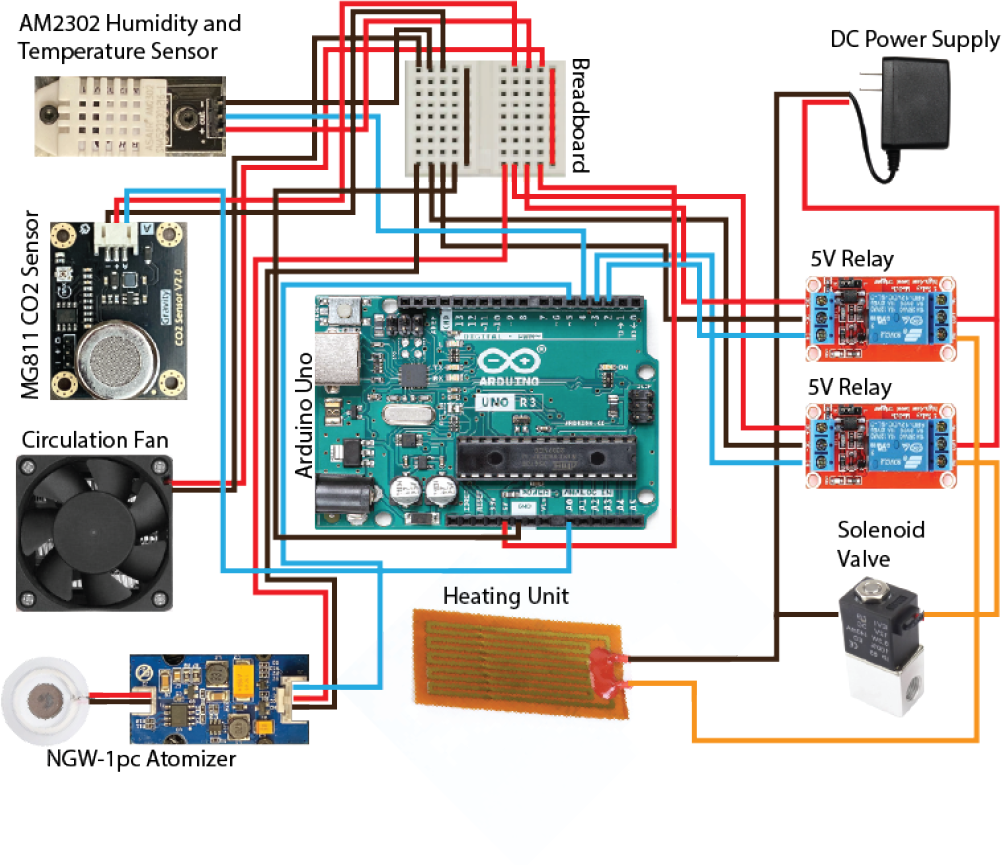
Circuit diagram of the device. Red and orange wires: positive connections (before and after passing through relays); black wires: negative connections; blue wires: signal connections

**FIG. 3.**
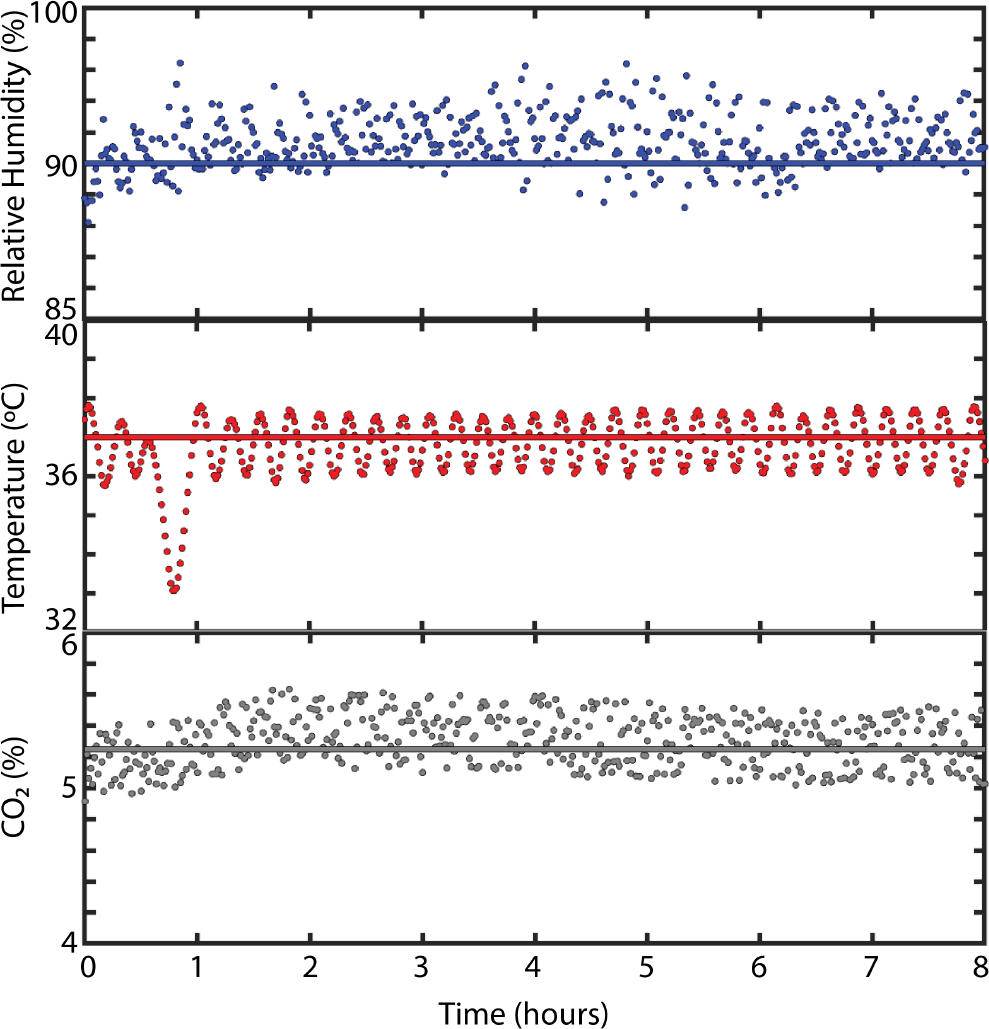
Environmental parameters inside the DIYncubator measured at one minute intervals over eight hours. At *t* = 0 the device had been running for 1 hour to come up to temperature, at which point humidity and CO_2_ control were engaged. (Top) Percent relative humidity; solid line indicates the set point of 90%, with mean humidity of 90.8 (Middle) Internal temperature, with set point of 37.0 °C, and mean temperature of 36.8 0.7 °C. (Bottom) Internal CO_2_ content, with set point of 5.3%, with equilibrium mean CO_2_ content of 5.3 *±* 0.2%

### B. Human Cell Growth Within the DIYncubator Compares Favorably to Commercial Solutions

To test the suitability of the device for live cell culture and imaging, we grew HEK293T cells in a 29 mm circular dish for 24 hours in the DIYncubator and compared the doubling time to the same cell line grown with identical conditions in a commercially purchased ibidi Blue Silver stage-top incubator (Fig. 4). The doubling time of HEK293T cells inside the DIYncubator was determined to be 39 *±* 6.5 hours, compared to a doubling time of 36 *±* 7.5 hours in the ibidi Blue Silver incubator.

**FIG. 4.**
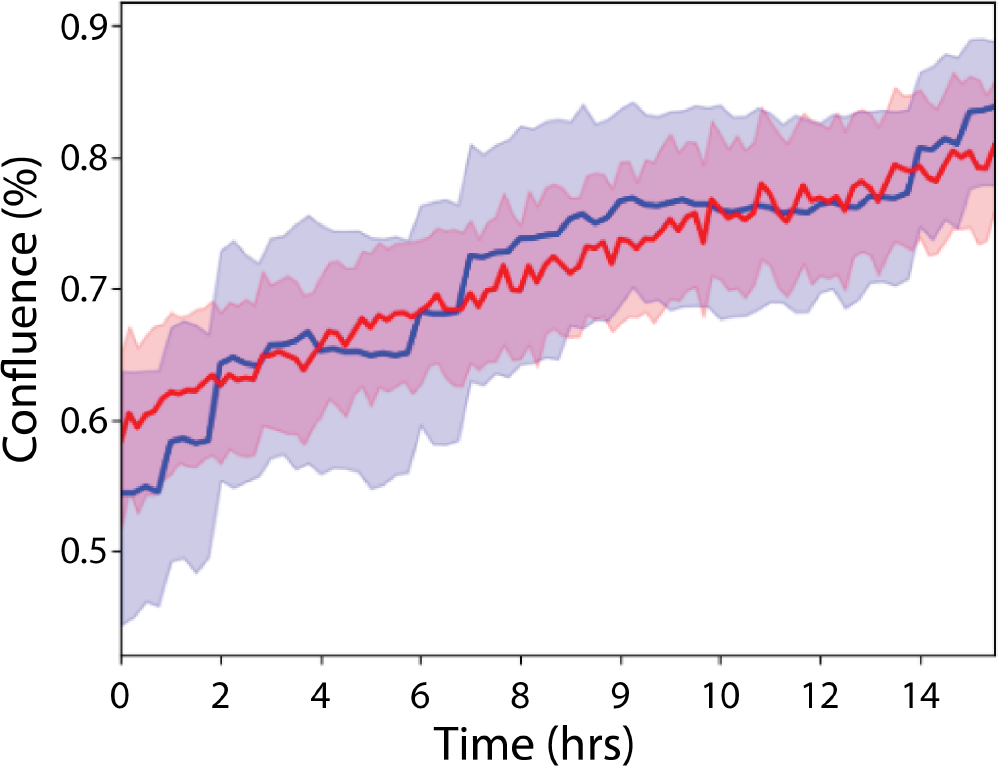
Comparison of growth rates between HEK293T cells cultured in an ibidi Silver Line stage-top incubator (blue) vs. the DIYncubator (red). Shaded regions indicate 95% confidence intervals, with solid lines indicating the mean over *∼* 200 cells.

## IV. DISCUSSION

We have described the design, construction, and implementation of a low cost, easy to build stage top incubation device called the “DIYncubator” for the growth and imaging of human and other mammalian cells. The DIYncubator is able to maintain an environment with controlled temperature, humidity, and CO_2_ content over long, multi-day time scales, allowing for fluorescent and brightfield imaging over the long times necessary to, for example, image oscillations in protein expression [8]. This performance compares favorable to commercially available solutions at *∼* 100x the cost.

However, with the advantage of the low cost comes a variety of disadvantages that should be kept in mind. The DIYncubator, as described here, was fabricated using ABS plastic, which is a low cost material but is relatively flexible and a poor conductor of heat. Additionally, care must be taken with heating approaches as ABS can warp considerably when exposed to high temperatures. Furthermore, the interior of the DIYcubator can become quite wet over long time periods, as the atomizer sprays moisture into the box and condensation forms on surfaces. As a result, the DIYcubator described here is ideal for fluorescence imaging where illumination is provided through the objective under the sample, but bright-field imaging accomplished by illumination through the removable lid may suffer somewhat as a result of the accumulation of condensation. However, the lid is easily removable and can periodically be wiped clean with minimal perturbation to the sample. Accumulation of condensation and moisture may also wreak havoc with electronic components over long times. However, after long periods of use we have not had significant failures of electronics, and, in any case, the electronics can easily be replaced many times over before the cost even begins to approach a fraction of that of commercially available systems.

As an alternative, the DIYcubator could be machined out of aluminum, which would yield a number of advantages. First, aluminum is more rigid, and would aid in creating a more airtight environment that would aid in reducing costs of operation by reducing the amount of CO_2_ required for a constant environment. Furthermore, the high thermal conductivity of aluminum could be utilized such that, by adhering the heating film directly to the chassis in a variety of locations, the entire body of the box could be used as a heating element to provide a more stable and homogeneous temperature profile across the unit. Finally, heating the chassis of the box in this way, and potentially combined with transparent heating film to make the removable lid, would substantially reduce condensation and therefore provide an environment more amenable to long-term brightfield illumination and imaging. However, acquiring and machining the requisite materials for this approach would add significantly to the financial and time cost, as well as the skills required, to fabricate the DIYncubator, the suitability of which is left to the discretion of the user.

